# Dynamics of microbial competition, commensalism and cooperation and its implications for coculture and microbiome engineering

**DOI:** 10.1101/2020.03.05.979435

**Authors:** Peng Xu

## Abstract

Microbial consortium is a complex adaptive system with higher order dynamic characteristics that are not present by individual members. To accurately predict the social interactions, we formulate a set of unstructured kinetic models to quantitatively capture the dynamic interactions of multiple microbial species. By introducing an interaction coefficient, we analytically derived the steady state solutions for the interacting species and the substrate profile in the chemostat. We analyzed the stability of the possible co-existing states defined by competition, parasitism, amensalism, commensalism and cooperation. Our model predicts that only parasitism, commensalism and cooperation could lead to stable co-existing state. We also determined the optimal social interaction criteria of microbial coculture with sequential metabolic reactions compartmentalized into two distinct species. Coupled with Luedeking–Piret and Michaelis-Menten equations, accumulation of metabolic intermediates in one species and formation of end-product in another species could be derived and assessed. We discovered that parasitism consortia disfavor the bioconversion of intermediate to final product; and commensalism consortia could efficiently convert metabolic intermediates to final product and maintain metabolic homeostasis with a broad range of operational conditions (i.e., dilution rates); whereas cooperative consortia leads to highly nonlinear pattern of precursor accumulation and end-product formation. The underlying dynamics and emergent properties of microbial consortia may provide critical knowledge for us to engineer efficient bioconversion process, deliver effective gut therapeutics as well as elucidate probiotic-pathogen interactions in general.

## 1. Introduction

Microbes in nature form diverse social interactions and dynamically respond to metabolic and environmental cues at community level. The interacting species in a microbial community might compete for the same resource, exchange for metabolites, communicate each other via metabolic or genetic signals (Fredrickson & Stephanopoulos, 1981). The unique interactions in a microbial community define the collective biological functions that are robust to harsh conditions when individual cells could hardly sustain (Brenner, You, & Arnold, 2008; Tsoi, Dai, & You, 2019). Compared to monocultures, co-cultures exhibit a number of advantages, including division of labor, compartmentalization of incompatible reactions, and robustness to perturbations et al (Jawed, Yazdani, & Koffas, 2019; McCarty & Ledesma-Amaro, 2018). Recently, microbial coculture or consortia has been increasingly applied to configurate different sections of metabolic pathways with improved catalytic performance. The co-cultivation of various species has led to the production of important biofuels (Minty et al., 2013), traditional food (Lu et al., 2018) and nutraceuticals (Arora et al., 2020; Wang, Zhao, Wang, & Koffas, 2020; Xu, Marsafari, Zha, & Koffas, 2020; Zhang & Wang, 2016). Novel synthetic biology tools, including cell signaling translator (Marsafari, Ma, Koffas, & Xu, 2020; Stephens, Pozo, Tsao, Hauk, & Bentley, 2019) and transcriptional factor-based biosensors (Lv, Gu, Xu, Zhou, & Xu, 2020; Lv et al., 2019; Rugbjerg, Sarup-Lytzen, Nagy, & Sommer, 2018) have been recently developed to autonomously regulate culture composition and eliminate metabolic heterogeneity. In particular, microbial social interaction could define unique spatial patterns that are important for us to fabricate advanced biomaterials (Ben Said, Tecon, Borer, & Or, 2020; Dai et al., 2019), understand biofilm formation (Beaudoin et al., 2017) and disarm antibiotic resistant superbugs (Davies & Davies, 2010).

Kinetic models have been increasingly important to help us understand microbial social interactions at the consortia-level (Kong, Meldgin, Collins, & Lu, 2018; Song, Cannon, Beliaev, & Konopka, 2014; Succurro & Ebenhöh, 2018). Most of these kinetic equations are developed by correlating cell growth fitness with the nutrient or environmental conditions of the interacting species. In particular, the canonical Monod equation, which describes the quantitative relation between cell growth and a limiting nutrient (Monod, 1949), has been expanded to incorporate multiple inhibitory terms (Han & Levenspiel, 1988; Levenspiel, 1980; Luong, 1987). For a 2-species coculture system, biochemical engineers have formulated a set of coupled Monod equations to describe the oscillatory relationship between Dictyostelium discoideum and Escherichia coli in Chemostat (Tsuchiya, Drake, Jost, & Fredrickson, 1972). On the other hand, ecologists preferred to use Logistic equation due to the simplicity, existence of analytical solution and the rich dynamics. For example, the solution of the Lotka–Volterra predator-prey model was derived and analyzed to describe the dynamic species interaction in a closed system (Lotka, 1926; Volterra, 1926). A recent hybrid Monod and Logistic model has been developed and solved to incorporate both nutrient-limiting conditions and self-inhibitory factors that may accurately describe cell growth (Xu, 2019).

Mathematical models of microbial consortia (Stephanopoulos, 1981) has been studied and analyzed in 1980s, which lay the foundation for us to understand microbial social interactions. However, the theoretical development of microbial consortia is not moving forward, partly due to the complex dynamics arising from the interacting species (Kong et al., 2018). Here we developed a set of unstructured kinetic models to quantitatively capture the dynamic interactions of multiple microbial species. We analytically derived the steady state solutions for the two interacting species and the substrate profile in the chemostat. By defining an interaction coefficient, we analyzed the stability of the possible co-existing states on the basis of eight microbial social interactions: competition, coexisting parasitism, extinctive parasitism, cooperation, bistable amensalism, extinctive amensalism, coexisting commensalism and extinctive commensalism. By analyzing the solutions for microbial consortia with sequential metabolic reactions compartmentalized into distinct species, we revealed the design criteria of microbial coculture engineering in chemostat. We discovered that commensalism consortia could efficiently convert metabolic intermediate to final product and maintain metabolic homeostasis (i.e., constant final product formation) with a broad range of operational conditions (dilution rates). The simplicity and the rich dynamics of the consortia model highlight the importance to incorporate social interaction parameters into the unstructured kinetic models. The dynamics of microbial competition and cooperation may facilitate us to assemble diverse microbial species with defined social interactions for important biotechnological and biomedical applications.

## 2. Computational methods

### 2.1 Matlab computational environment

Matlab R2017b was used as the computational platform and installed on a Windows 7 professional operation system with Intel Core i3-6100 CPU processor at speed of 3.70 GHz. The installed memory (RAM) is 4.0 GHz. Matlab symbolic language package coupled with LaTex makeup language is used to derive and output the symbolic equations and solutions (Supplementary files). Analytical solutions were derived for Fig. 2 and Fig. 4 with Matlab Symbolic Language. All trajectories in Fig. 2 and Fig. 4, and phase portraits in Fig. 3 were computed by numerical ODE45 solvers. Numerical solutions for the compartmentalized sequential metabolic reactions in Fig. 5 and Fig. 6 were computed by numerical ODE45 solver. Matlab code with symbolic functions and m.files has been compiled into the supplementary file.

**Fig. 1.**
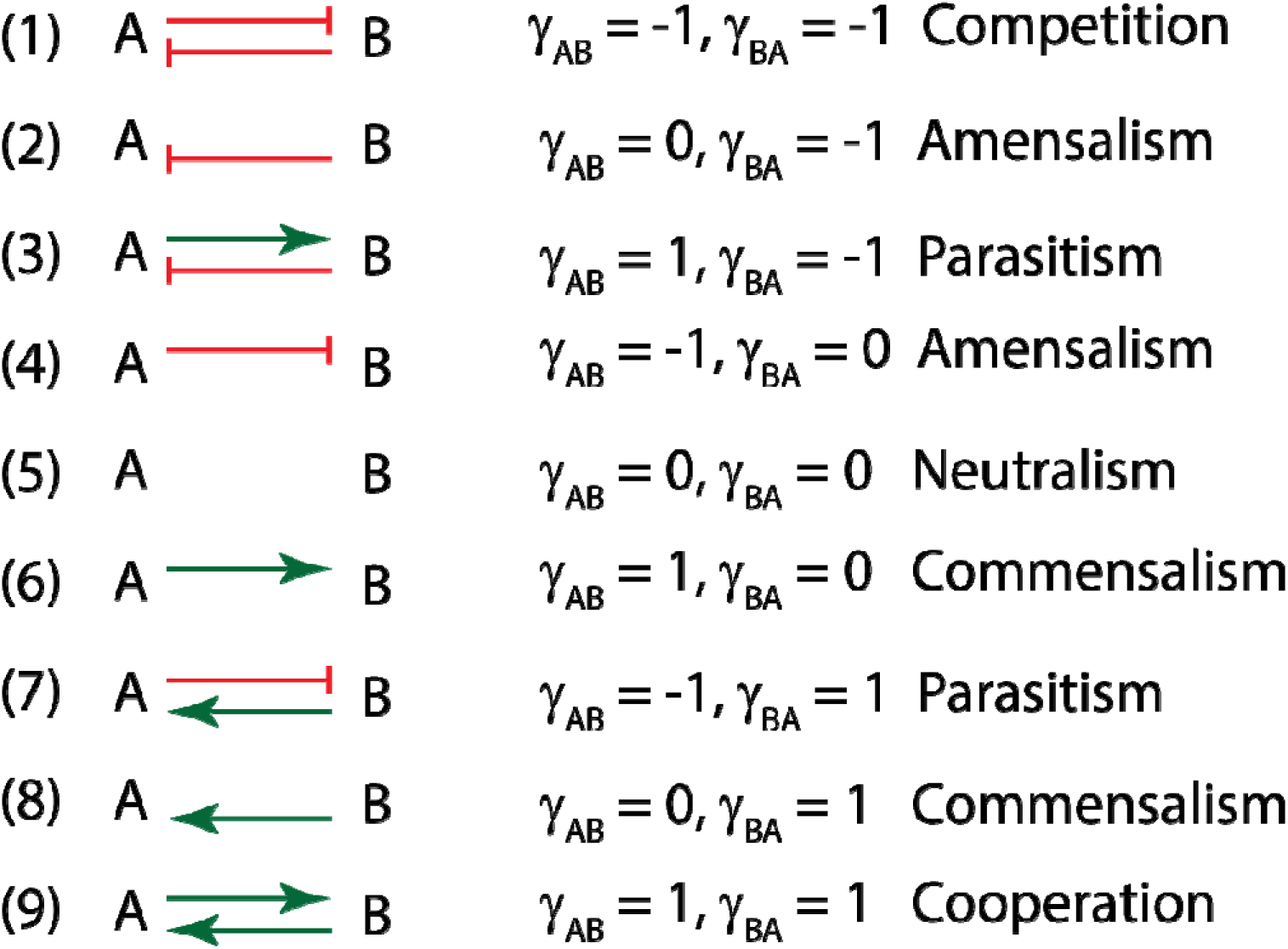
Categories of microbial social interactions: competition, amensalism, parasitism, neutralism, commensalism and cooperation. The interaction coefficient was defined by a dimensionless factor (γ_AB_ *or* γ_BA_) that describe the beneficial or detrimental interactions between species A and species B. A green arrow indicates beneficial relation, a blunt-ended orange arrow indicates harmful relation.

**Fig. 2.**
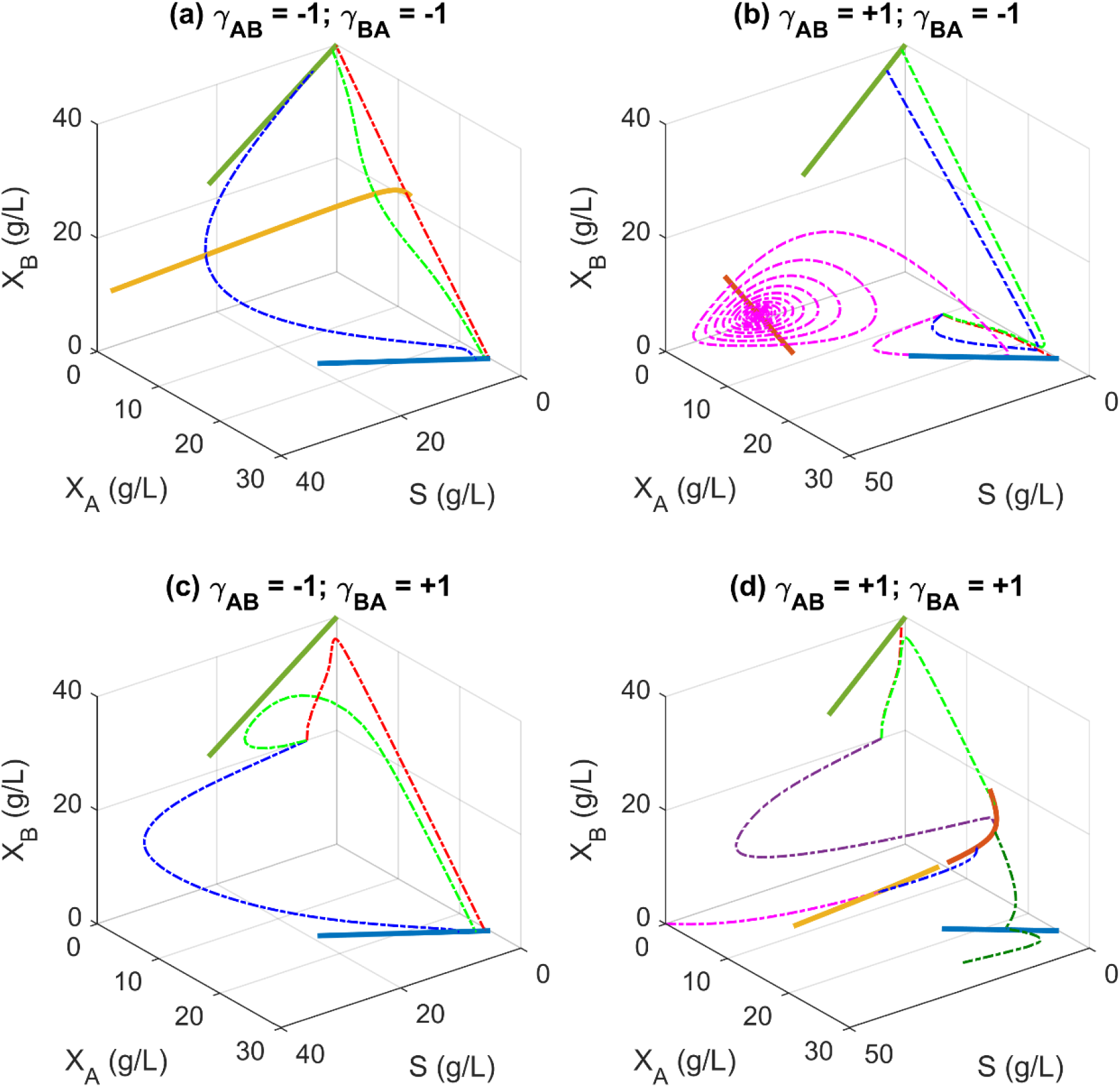
Dynamics of microbial competition, parasitism and cooperation at different dilution rate in chemostat. Green solid line: species B exist alone; Blue solid line: species A exists alone; Light orange solid line: *unstable* co-exist solution; Dark orange solid line: *stable* co-exist solution; Dash line: trajectory of steady state solution due to a small perturbation from any random state with fixed dilution rate. In all the simulation, we assume that species A has a larger growth fitness (maximal specific growth rate) than species B. (**a**) Competition. Species A and species B mutually exclude each other, leading to an unstable co-exist solution. Any perturbation from the coexist state (light orange line) will result in the survival of a single species (either A or B). (**b**) Coexisting parasitism: Species A benefits species B, but species B is harmful to species A. Stable coexisting is possible at relatively large dilution rate (equivalently to harsh conditions). (**c**) Extinctive parasitism: Species B benefits species A, but species A is harmful to species B. Stable coexisting is impossible and B will extinct. (**d**) Cooperation: Species A and species B mutually benefit each other. A stable coexisting solution (dark orange line) is possible due to the mutualistic interactions between species A and B. Any infinitesimal perturbation from the unstable coexisting solution (light orange line) will move the system to a washout state (solution falls to the origin) or to the stable co-existing state (dark orange line).

**Fig. 3.**
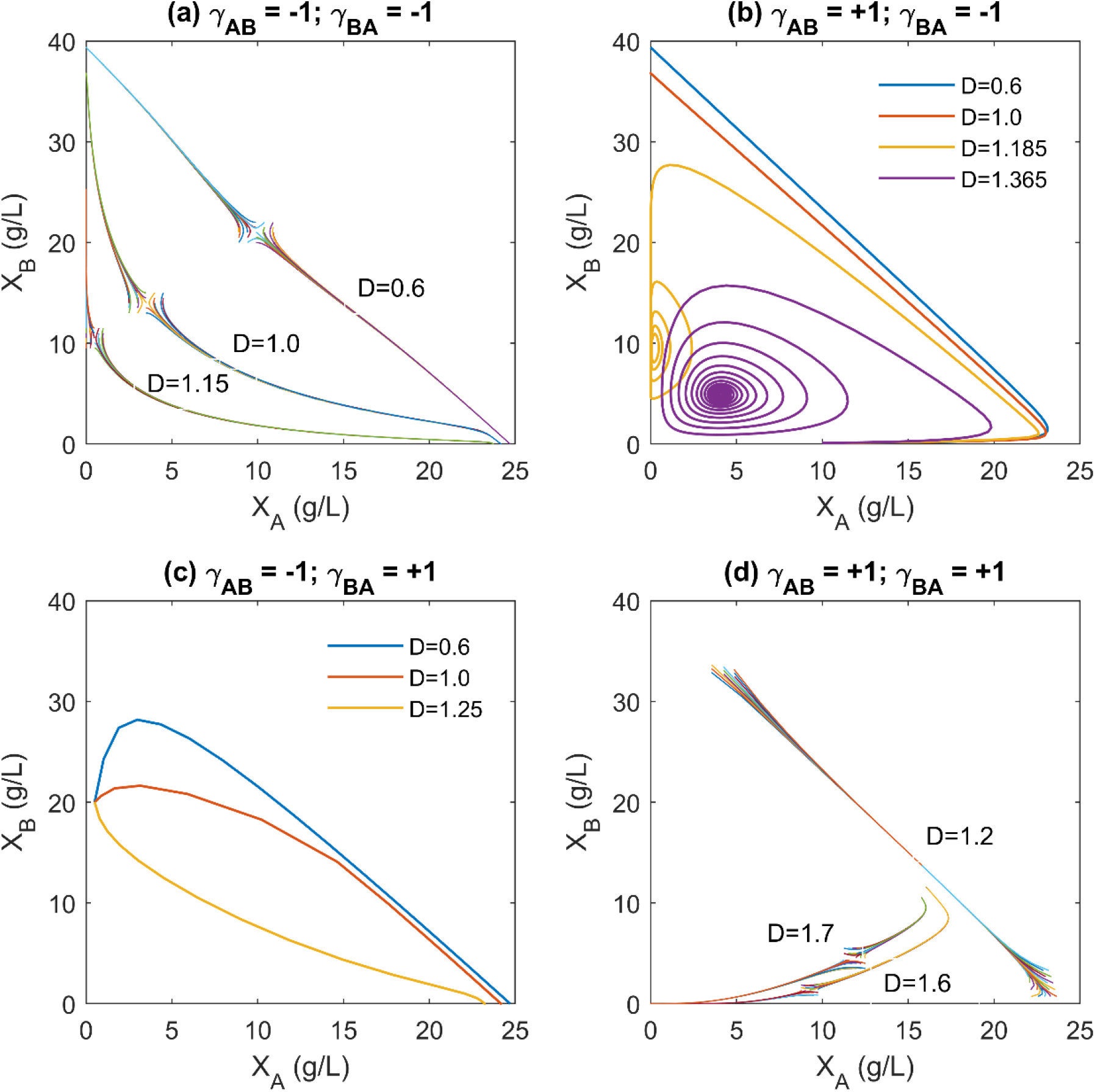
Two-dimensional phase portrait of steady state solutions of simple microbial interaction models. Phase portrait for bistable competition (**a**), coexisting parasitism (**b**), extinctive parasitism (**c**), and cooperation (**d**). The 2-D phase portrait corresponds to the trajectory of the solution from any random state at a fixed dilution rate in Fig. 2. In all the simulation, we assume that A has a larger growth fitness than species B.

**Fig. 4.**
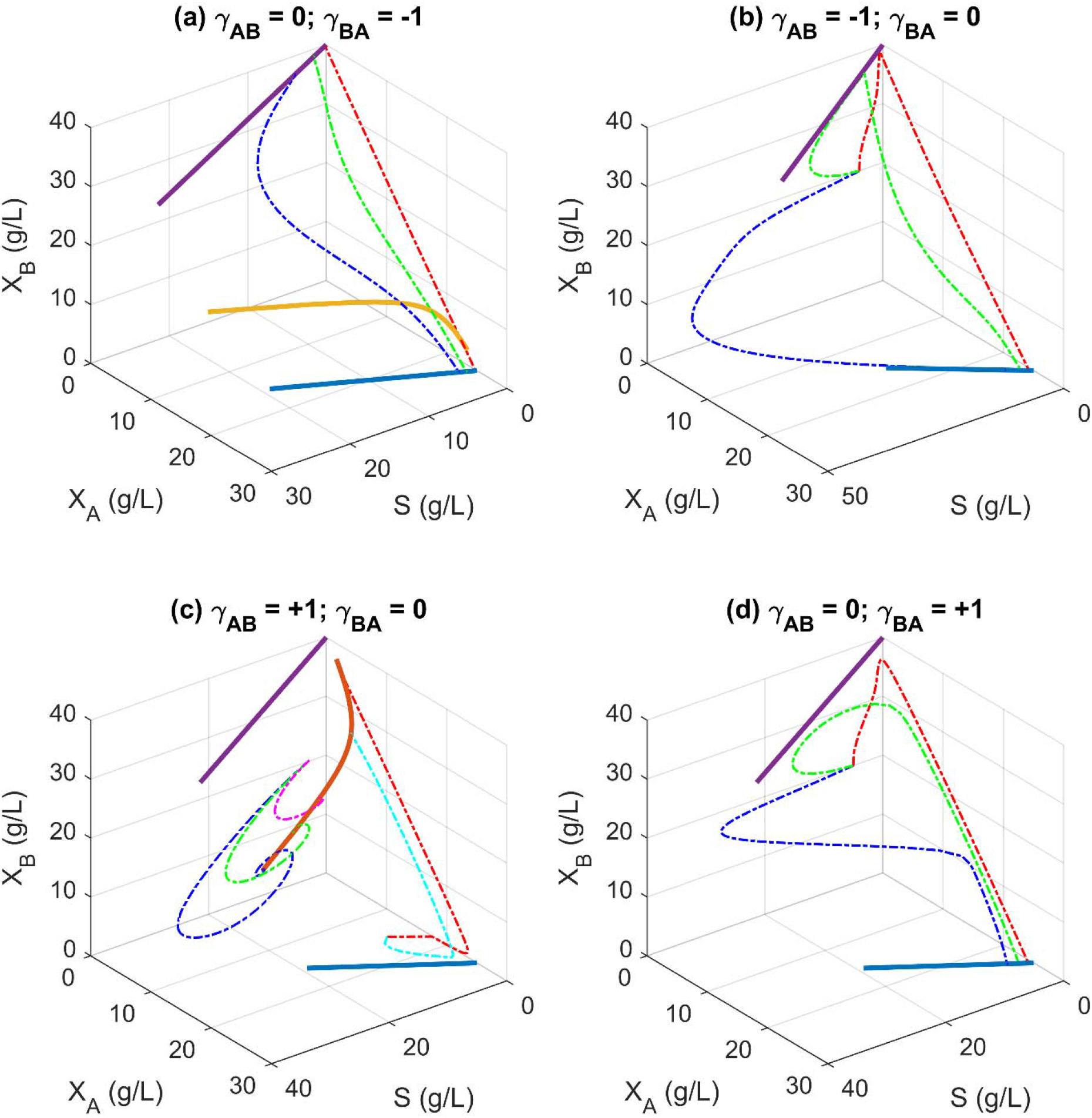
Dynamics of microbial amensalism and commensalism at different dilution rate in chemostat. Purple solid line: species B exist alone; Blue solid line: species A exists alone; Light orange solid line: *unstable* co-exist solution; Dark orange solid line: *stable* co-exist solution; Dash line: trajectory of steady state solution due to a small perturbation from any random state with fixed dilution rate. In all the simulation, we assume that species A has a larger growth fitness than species B. (**a**) Bistable amensalism: Species B is harmful to species A, but species A is neutral to species B. Coexistence is unstable and any perturbation will lead to the survival of a single species (either species A or species B). (**b**) Extinctive amensalism: Species A is harmful to species B, but species B is neutral to species A. The system eventually leads to the existence of species A alone (species B will extinct). (**c**) Coexisting commensalism: Species A benefits species B, but species B is neutral to species A. Species A and B will exist together. (**d**) Extinctive commensalism: Species B benefits species A, but species A is neutral to species B. The system eventually leads to the existence of species A alone (species B will extinct).

**Fig. 5.**
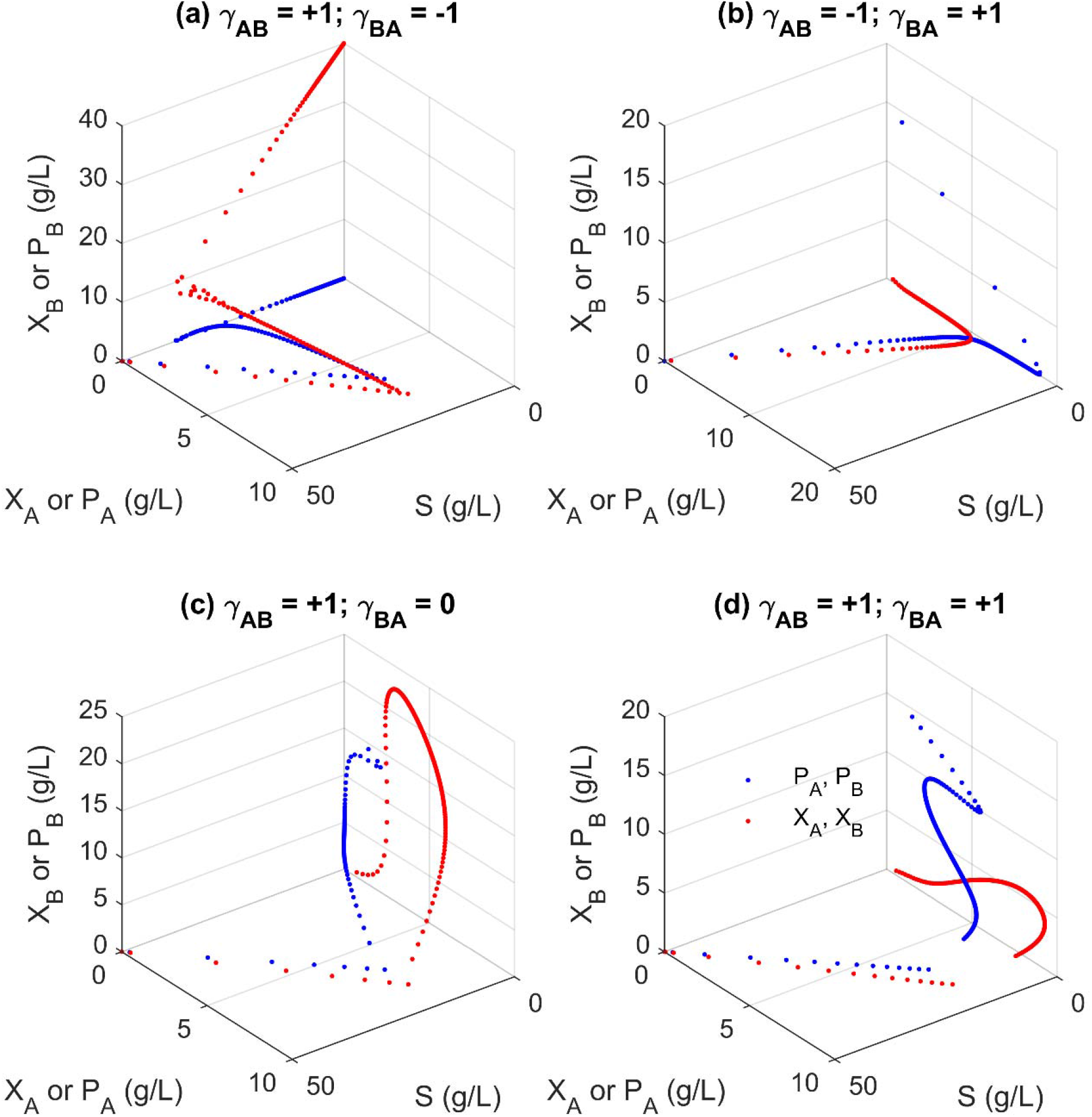
Steady state solutions of microbial co-culture with sequential metabolic reactions compartmentalized in two species. Red dots: biomass; blue dots: intermediate *P*_A_ or product *P*_B_. In all the simulation, we assume that species A has a larger growth fitness than species B. Intermediate *P*_A_ secreted from species A is converted to product *P*_B_ by species B. (**a**) Coexisting parasitism: Species A benefits species B, but species B is harmful to species A. (**b**) Extinctive parasitism: Species B benefits species A, but species A is harmful to species B. The system will move to species A existing state with only intermediate *P*_A_ accumulation (no product *P*_B_ formation). (**c**) Coexisting commensalism: Species A benefits species B, but B is neutral to A. (**d**). Cooperation: Species A and B mutually benefit each other.

**Fig. 6.**
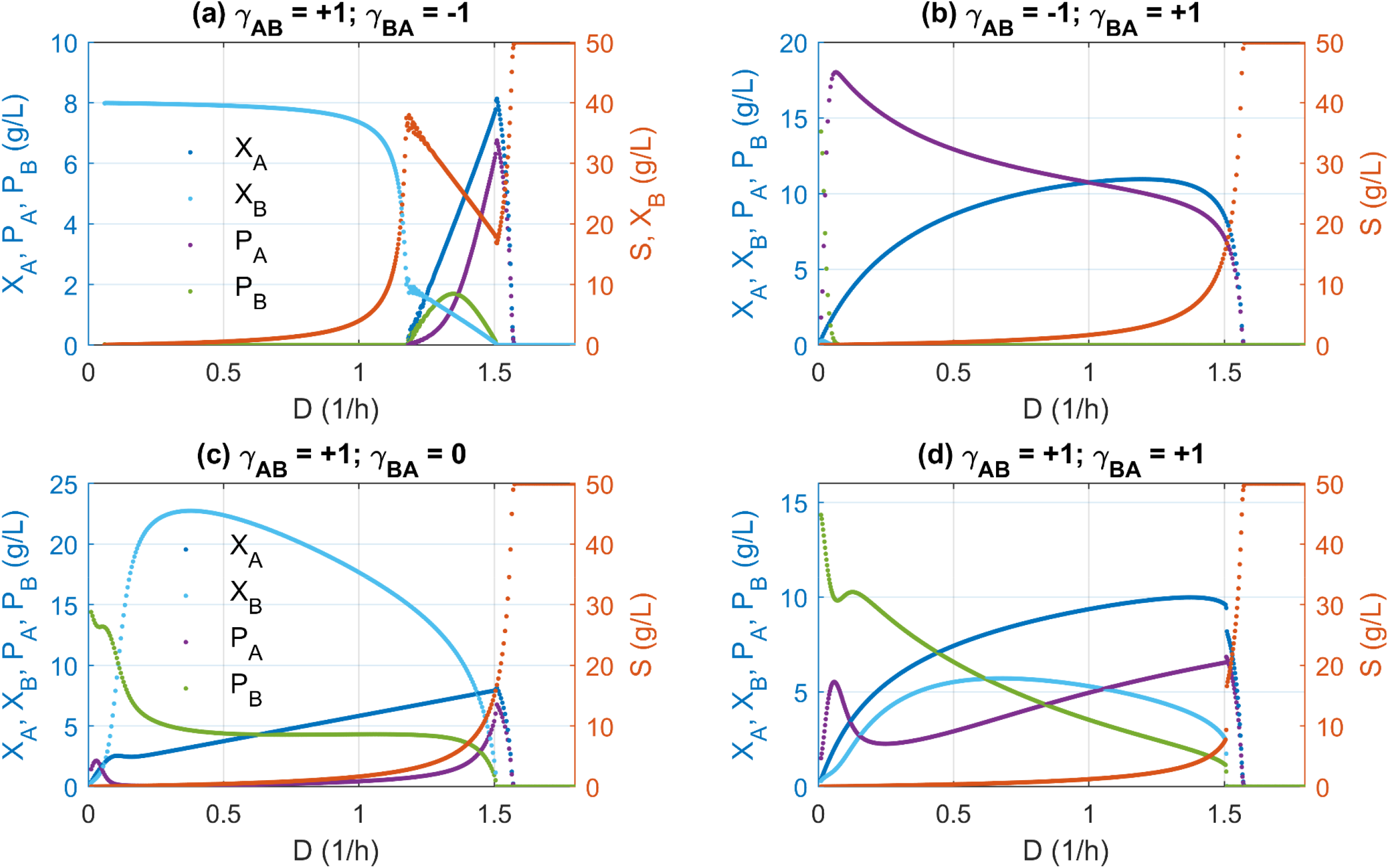
Operational conditions of microbial co-culture with sequential metabolic reactions compartmentalized in two species. (**a**) Coexisting parasitism leads to the inefficient conversion of intermediate *P*_A_ to product *P*_B_, as the dilution rate increases. Optimal dilution rate is possible to maximize *P*_B_ production. (**b**) Extinctive parasitism only leads to the presence of species A and the accumulation of intermediate PA. Optimal dilution rate is possible to maximize intermediate *P*_A_ production. (**c**) Coexisting commensalism leads to the coexistence of species A and B and the efficient conversion of intermediate *P*_A_ to product *P*_B_. (**d**) Mutualistic cooperation leads to the accumulation of intermediate *P*_A_ and rapid declining of product *P*_B_ as the dilution rate increases.

### 2.2 Stability analysis of microbial social interactions

For the generalized systems ODE equations listed in Eqn. 3, Eqn. 4 and Eqn. 5, we have computed the parameterized Jacobian matrix with Matlab symbolic language. This parameterized Jacobina matrix represents all the nine social interactions summarized in **Fig. 1**. One can simply substitute specific numbers (−1, 0, or 1) into the interaction coefficients (γ_AB_ or γ_BA_) in this generalized Jacobian matrix to retrieve the detailed matrix. Then specific eigenvalues cold be computed to evaluate the stability criteria of each steady states. A complete long form of the Jacobian Matrix is provided as supplementary equations. Some representative eigenvalues for specific parameter conditions are compiled to the supplementary table. Here is the generalized Jacobian Matrix that corresponds to systems ODEs Eqn.3 to Eqn. 5.

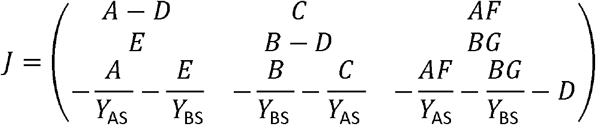

Where *A, B, C, D, E, F* and *G* represents

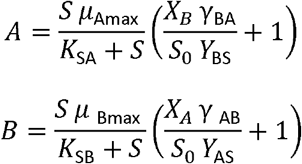

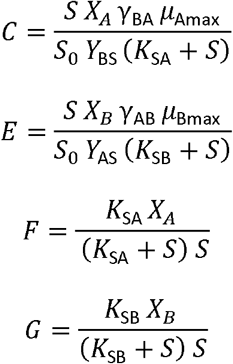

### 2.3 Parameter settings

All the solutions were derived either analytically or numerically with the following parameter settings: *μ*_Amax_= 1.6 (1/h); *μ*_Bmax_ = 1.2 (1/h); *K*_SA_ = 1.0 (*g/L*); *K*_SB_ = 0.8 (g/L); *S* = 50(*g/L*); *Y*_AS_ = 0.5 (g/g); *Y*_BS_= 0.8 (g/g); *Y*_BA_ = 0.8 (g/g); *Y*_PS_ = 0.4 (g/g); *α* = 0.5 (dimensionless); *β* = 0.5 (1/h); *γ*_AB_ = −1, 0, or 1 (dimensionless), depending on the social interactions as specified in **Fig. 1**; *γ*_BA_ = −1, 0, or 1 (dimensionless), depending on the social interactions as specified in **Fig. 1**; %= 0.8 (1/h); *K*_*m*_ = 1.0 (g/L). Dilution rate *D* could be varied from 0 to 1.8 (1/h). All biophysical parameters were taken from biochemical parameter database (BioNumbers) or biochemical engineering textbooks (Shuler, Kargi, & DeLisa, 2017) with a physiologically interpretable range. It should be noted that, we assume that species A has a larger growth fitness (maximal specific growth rate) than species B in all our simulations.

## 3. Results

### 3.1 A unstructured kinetic model to define microbial social interactions

Based on the beneficial or detrimental relationship between two species, we can define six social interactions (**Fig. 1**): competition, amensalism, parasitism, neutralism, commensalism and mutualism (cooperation). To define the dynamic nature of microbial consortia, we modified the growth fitness function (**Eqn.1** and **Eqn. 2**) by introducing an interaction coefficient (γ_AB_ or γ_BA_) to each of the interacting species. An interaction factor (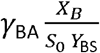 or 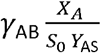), which is defined as the product of the interaction coefficient and relative population of the interacting species, is incorporated to the growth fitness equation as specified by Eqn.1 and Eqn.2.

The benefits of introducing this interaction coefficient are multifold. For example, the growth fitness equation (**Eqn.1** or **Eqn. 2**) converges to the canonical Monod equation, when the population of the interacting species is negligible (*X*_*A*_ → 0 or *X*_*B*_ → 0). As the population of the interacting species approaches to the capacity of the system (*X*_*A*_ → *S*_0_ *Y*_*AS*_ or *X*_*B*_ → *S*_0_ *Y*_*BS*_), the interaction factor (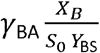 or 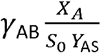) reaches its maximal or minimal value (γ_BA_ or γ_AB_) that corresponds to either beneficial or detrimental effects, depending on the sign (+1 or −1) of the interaction coefficients (γ_AB_ or γ_BA_). As the cell grows, the population starts having beneficial or detrimental effect on the interacting species; and the strength of this interaction is proportional to the relative ratio of the cell populations. The sign (negative or positive) of the interaction coefficient (γ_AB_ or γ_BA_) determines the nature (detrimental or beneficial) of the social interactions (**Fig. 1**). Following classical biochemical kinetics, we formulated a set of simplified microbial consortia models (Eqn. 1 to Eqn. 8). With these equations, we will derive the analytical solutions and evaluate the stability of the co-existing states. The insight obtained from the model will be critical for us to understand the dynamic nature of microbial consortia and provide us computational guidelines to maintain stable microbial coculture, which may facilitate us to design efficient microbial process for biomanufacturing or gut therapeutics with human health benefits.

### 3.2 Dynamics of microbial competition, parasitism and cooperation with resource limitations

We next sought to understand the dynamics of microbial consortia with strong interactions, namely competition, parasitism and cooperation. One important question we would like to answer is whether the two interacting species could stably co-exist. This is critical, because there would be no microbial consortia if the two species could not grow together. With Matlab Symbolic computation package, we analytically derived the steady state solutions for Eqns. 1 to Eqns. 5 (a supplementary Matlab code was provided as SI file). By varying the interaction coefficients (γ_AB_ or γ_BA_), we could use the set of parameterized equations (Eqn. 1 to Eqn. 5) to describe the various social interactions of the microbial consortia.

When species A and species B mutually exclude each other and compete for the same substrate, this condition γ_AB_ = −1 γ_BA_ = −1) will lead to an unstable co-existing state (**Fig. 2a**), as evidenced that one of the eigenvalues of Jacobian Matrix is positive (Supplementary Eigenvalue tables). Indeed, any perturbation from the co-existing state (light orange line in **Fig. 2a**) will make the system traverse to a single species survival state (**Fig. 3a**), without the presence of the other species. The two species mutually exclude each other; therefore they could not grow together in CSTR. For example, as we increase the dilution rate (*D* = 0.2 for red trajectory, *D* = 0.6 for green trajectory and *D* = 1.0 for blue trajectory in **Fig. 2a**), the unstable coexisting state will traverse toward either species A or species B existing state (**Fig.2a** and **Fig. 3a**). The steady state solution of species A or B decreases as we increase the dilution rate. This trend is consistent with the population concentration as observed when a single species is cultivated in the chemostat (Xu, 2019).

When species A benefits species B, but species B harms species A, this condition γ_AB_ = +1 *and* γ_BA_ = −1) will lead to a coexisting parasitism regime (**Fig. 2b**). At relatively low dilution rate (*D* < 1.185 h^−1^), species B will win and dominate regardless of the initial concentration of species A (**Fig. 2b** and **Fig. 3b**). For example, the trajectory of the solution will move to species B alone state, as we increase the dilution rate (*D* = 0.6 for green trajectory and *D* = 1.0 for blue trajectory in **Fig. 2a**). Interestingly, at relatively large dilution rate (*D* ≥ 1.185 h^−1^), the system will move to a parasitism coexisting state (dark orange line in **Fig. 2b**). Since this dilution rate is approaching to the maximal growth rate of species B (*μ*_Bmax_ = 1.2 h^−1^), species B must depend on the existence of species A to gain growth advantage. For example, species A may produce a public good that benefits the growth of species B. As a result, we observed an oscillatory trajectory (a limited cycle, **Fig. 2b** and **Fig. 3b**) where species A and species B eventually lead to a balanced distribution (*D* = 1.365), as specified by the dark orange line in **Fig. 2b**. This parasitism coexisting state is stable since all the eigenvalues are negative (Supplementary Eigenvalue tables). On the contrary, if we flip the sign of the interaction coefficient *γ*_AB_ = −1 *γ*_BA_ = +1), the system will lead to an extinctive parasitism: where that species A has a large growth fitness than species B (*μ*_Amax_ > *μ*_Bmax_). This analysis indicates species A will always outcompete the growth of species B (**Fig. 2c** and **Fig. 3c**), due to the fact that parasitism interaction could either lead to co-existing or extinction sates. The exploitative relationship between species A and species B may reach a coexisting state, only if the dilution rate is harmful to the exploiter (i.e., species B is an exploiter in Fig. **2b**).

When species A and B mutually benefit each other, this condition *γ*_AB_= −1 γ_BA_ *and* = −1) will lead to a cooperative state (**Fig. 2d**). A stable coexisting state (dark orange line) is possible due to the mutualistic interactions between species A and B. Interestingly, when the dilution rate is approaching to the maximal growth rate of species A (*µ*_Amax_ = 1.6 h^−1^), the system will move to a unstable co-existing state (the light orange line in Fig. **2d**). Any infinitesimal perturbation from this unstable coexisting solution (light orange line) will move the system to an extinction state (i.e., solution falls to the origin, pink trajectory for *D* = 1.7) or to the stable co-existing state (i.e., solution falls to the dark orange line, blue trajectory for *D* = 1.7). This intriguing bifurcating behavior is also exemplified in the 2-D phase portrait as *D* is above 1.58 (Fig. **3d**). This bistability at large dilution rate (*D* > 1.58) suggests that the relative population of species A and B is critical to maintain the co-existing state. Obviously, the cooperation allows the system to operate at a dilution rate that is larger than both species could sustain, indicating the robust nature of the mutualistic interactions.

### 3.3 Dynamics of microbial commensalism and amensalism with resource limitations

We next sought to understand the dynamics of microbial consortia with weak interactions, namely amensalism and commensalism. Under these conditions, we are also interested in understanding whether the two interacting species could co-exist stably or not. Following the same stability criteria, we evaluated the eigenvalues of Jacobian matrix. According to the system dynamics behavior, we could categorize the weak interactions into four groups: bistable amensalism (**Fig. 4a**), extinctive amensalism (**Fig. 4b**), coexisting commensalism (**Fig. 4c**) and extinctive commensalism (**Fig. 4d**).

When species B is harmful to species A, but species A is neutral to species B, the system (*γ*_AB_ = 0 *and γ*_BA_ = 1) will lead to an unstable coexisting state (light orange line of **Fig. 4a**). For example, any perturbation from this coexisting state will lead to the survival of a single species (either species A or species B). As we increase the dilution rate (*D* = 0.2 for red trajectory, *D* = 0.8 for green trajectory and *D* = 1.0 for blue trajectory in **Fig. 4a**), the unstable coexisting state will traverse toward either species A or species B existing state (**Fig. 4a**). We named this as “Bistable amensalism”. On the contrary, an extinctive amensalism will arise when species A is harmful to species B, but species B is neutral to A (*γ*_AB_ = −1 *and γ*_BA_ = 0). Under any dilution condition, species A will outcompete species B to exist alone in the system (**Fig. 4b**), due to the fact that species A has a large growth fitness than species B (*μ*_Amax_ > *μ*_Bmax_).

When species A benefits species B, but species B is neutral to species A, the system (*γ*_AB_ = +1 *and γ*_BA_ = 0) will move to a stable coexisting state (dark orange line of **Fig. 4c**). For example, with a small perturbation from any initial conditions, the trajectory of the system solution will traverse toward the co-existing state (the dark orange line of **Fig. 4c**). Under this condition, the species with larger growth fitness (i.e., species A) will promote the growth of the species with smaller growth fitness (i.e., species B). On the contrary, an extinctive commensalism will arise if species A is neutral to species B, but species B benefits species A (*γ*_AB_ = 0 *and γ*_BA_ = +1). Under any dilution condition, species A will outcompete species B to exist alone in the system (**Fig. 4d**).

In summary, the social interactions of two species with resource limitations could result in rich dynamics ranging from bistable competition (**Fig. 2a**), coexisting parasitism (**Fig. 2b**), extinctive parasitism (**Fig. 2c**), cooperation (**Fig. 2d**), bistable amensalism (**Fig. 4a**), extinctive amensalism (**Fig. 4b**), coexisting commensalism (**Fig. 4c**) and extinctive commensalism (**Fig. 4d**). This complex interaction was captured by a set of unstructured kinetic models developed in this work, simply with the introduction of the interaction coefficients (*γ*_AB_ *or γ*_BA_) to the classical Monod equations. The parameterized system equations (**Eqn.1** to **Eqn. 5**) provides a quantitative approach to analyze the system dynamics, which may help us design more efficient consortia and leverage coculture engineering for biotechnological and biomedical applications. It should be noted that neutralism (where two species grow independently) was not explored in this study, but neutralism was also captured in our model (*γ*_AB_ = 0 *and γ*_BA_ = 0, **Fig. 1**)

### 3.4 Implications for microbial co-culture engineering and microbiome engineering

Co-cultures or microbial consortia are emerging strategies to improve metabolic pathway efficiency. They exhibit a number of advantages over monoculture, including division of labor, compartmentalization of reaction, and robustness to perturbations (McCarty & Ledesma-Amaro, 2018; Wang et al., 2020; Zhang & Wang, 2016). Microbial consortia define the functional assembly and social interaction of multiple species. To design efficient bioconversion process, one must consider the social interactions of the individual members in the microbial community. In this section, we will determine the optimal social interaction criteria for microbial coculture when a 2-step sequential metabolic reaction is compartmentalized into two co-cultivating species. Namely, intermediate A (*P*_*A*_) is secreted from species A and was later converted to final product B (*P*_*B*_) by species B. Two additional equations (**Eqn. 6** and **Eqn. 7**) and a modified substrate consumption equation (**Eqn. 8**) were introduced to describe the system dynamics. We assume there is no bottleneck of metabolites transportation/diffusion across the two species and the consortia is devoid of metabolic burden due to accumulation of intermediate or final product. The formation rate for intermediate A (*P*_*A*_) was assumed to follow the Luedeking-Piret equation (Robert Luedeking, 1959), and the formation rate for product B (*P*_*A*_) should follow a Michaelis-Mention type kinetics with the turnover rate constant proportional to the concentration of species B. With biomass yield (*Y*_AS_ *and Y*_BS_), intermediate A yield from substrate (*Y*_PS_), and product B yield from intermediate A (*Y*_BA_), we can derive the mass balance equation for substrate consumption as specified by **Eqn. 8**.

To simplify the discussion, we focus on the steady state solutions of coexisting parasitism (**Fig. 5a** and **Fig. 6a**), extinctive parasitism (**Fig. 5b** and **Fig. 6b**), coexisting commensalism (**Fig. 5c** and **Fig. 6c**), and cooperation (**Fig. 5d** and **Fig. 6d**). The distribution of the steady state solution with varying dilution rate at 3-D space is presented in **Fig. 5**, and the exact steady state solution with varying dilution rate at a 2-D panel is presented in **Fig. 6**.

The solution distribution for biomass (*X*_*A*_ and *X*_*B*_) and metabolite (*P*_A_ and *P*_B_) displays highly nonlinear behavior (**Fig. 5** and **Fig. 6**), especially for the case when the two species coexist. Consistent with previous findings (**Fig. 2b**), coexisting parasitism (*γ*_AB_ = +1 *and γ*_BA_ = −1) allow species B to exist alone and species A is suppressed by species B at low dilution rate (*D* < 1.185 h^−1^). Therefore, intermediate A (*P*_*A*_) was produced at minimal value (~0) and there is value (*D* ≥ 1.185 h^−1^) in proximate to the maximal growth rate of species B (*μ*_Bmax_ = 1.2 h^−1^), no B produced at low dilution rate (**Fig. 5a** and **Fig. 6a**). When the dilution rate reaches a critical species B must rely on the beneficial effects (public goods or welfare) of species A to gain growth fitness (**Fig. 5a** and **Fig. 6a**). In other words, species A will be encouraged to proliferate, and the two species will coexist with a balanced population distribution. Under high dilution −1 rate (*D* ≥ 1.185 h^−1^), intermediate A (*P*_*A*_) secreted from species A will be converted to product B (*P*_B_) by species B (**Fig. 5a** and **Fig. 6a**). As predicted in the simulation, we can arrive an optimal dilution rate to maximize the product B formation (*P*_B_), despite the fact that there is a tipping (discontinuous) point for species A (*X*_A_) and intermediate A (*P*_*A*_) (**Fig. 6a**) as we increase the dilution rate. This optimal dilution rate could be analytically derived should we have enough computational power. On the contrary, an extinctive parasitism system (*γ*_AB_ = −1 *and γ*_BA_ = +1) will allow species A exist alone, hence only the accumulation of the intermediate A (*P*_*A*_) without formation of product B (**Fig. 5b** and **Fig. 6b**). The solution will eventually fall to the origin (washout state) when we further increase the dilution rate for all the scenario discussed here (**Fig. 5** and **Fig. 6**). This analysis indicates that a parasitism relationship may allow the compartmentalization of two sequential metabolic reactions in two distinct species, and there exists an optimal dilution rate to maximize the metabolite production (*P*_B_). In reality, this metabolite *P*_B_ might be related with some signaling molecules that are associated with antibiotic resistance in biofilm formation, secondary metabolite synthesis in endophytic fungi or a metabolic intermediate that is associated with the dysbiosis of gut microbes in living organisms.

The most complicated dynamics are displayed when the two species are commensal (**Fig. 5c**) or cooperative (**Fig. 5d**). In both scenario, coexisting is possible and product B will be formed when the two species harbor distinct sections of metabolic reactions (**Fig. 6c** and **Fig. 6d**). In particular, intermediate A (*P*_A_) secreted from species A will be efficiently converted to product B (*P*_B_) when the two specie form a commensalism consortium (**Fig. 5c** and **Fig. 6c**). As a result, the intermediate A (*P*_A_) was almost kept at minimal level with a very large window of operational conditions (i.e, 0.16 < *D* < 1.32), as evidenced that there is almost constant product B (*P*_B_) formed in the system (**Fig. 6c**). We can also arrive an optimal dilution rate to maximize biomass for species B (**Fig. 6c**). In addition, the simulation indicates drastic changes of species A biomass (*X*_A_) and intermediate (*P*_A_) at a tipping point when the dilution rate is about 1.5 h^−1^. Beyond this tipping point, the system rapidly falls to the washout states (**Fig. 5c** and **Fig. 6c**). As a comparison, cooperation between species A and B instead leads to the accumulation of intermediate A (*P*_A_ in **Fig. 5d** and **Fig. 6d**), which possibly due to the mutualistic beneficial interaction between species A and species B: increase in the biomass of species A, which is the source for intermediate *P*_A_, will benefit the growth fitness of species B, which is the sink for intermediate *P*_A_. Under this scenario, the activity of the metabolic source pathway and metabolic sink pathway is proportionally increased. Due to the intrinsic parameter settings (i.e., species A has a larger fitness than species B), intermediate A (*P*_A_) from the metabolic source strain might not be efficiently converted to product B by the metabolic sink strain. This might possibly explain the inefficient conversion of intermediate A (*P*_A_) to product B (*P*_B_). Biomass (*X*_A_) and intermediate (*P*_A_) in species A will keep increasing before the system reaches the tipping dilution rate (D ≈ 1.5 h^−1^) (**Fig. 6d**). Product B will keep decreasing within the operational −1 −1 dilution window (0 h^−1^ < D < 1.6 h^−1^). This analysis indicates that designing a cooperative consortium will be more challenging than designing a commensal consortium due to the mutualistic interaction of the two species. Importantly, commensal consortia allow the stable existence of two species and the efficient conversion of metabolic intermediate (*P*_A_) to final product (*P*_B_). In particular, metabolite concentration (*P*_B_) was almost kept constant at a very large dilution window (i.e, 0.16 < D < 1.32), which may explain the phenomenon why most of the species in gut microbiota maintain a commensal consortium. Equivalently to say, this commensal consortium is critical to maintain metabolite homeostasis (i.e. constant *P*_B_), which could resist large perturbations of environmental condition change (i.e., the dilution rate discussed in this study or food uptake/digestion rate in human gut).

## 4. Discussions

From a 2-strain Lotka–Volterra competition model, Ram et al have been able to predict the microbial growth fitness of individual species from the growth curve data of a mixed cell culture (Ram et al., 2019). Their model contains a number of biological factors that dictate cell growth fitness, including specific growth rate at low density, maximum cell density, deceleration parameter and a frequency-based adjustment function. By fitting the monoculture growth data to the Baranyi–Roberts model, the authors were able to retrieve these critical parameters. Remarkably, a competition coefficient was found sufficient to predict the growth behavior of the mixed cell populations. Later Balsa-Canto et al argued this approach may recover the steady states, but it may fail to reproduce the dynamics of the subpopulations in the mixed cell culture (Balsa-Canto, Alonso-del-Real, & Querol, 2020). Errors of the estimated competition parameters in the vicinity of the boundaries between coexistence and exclusion may lead to biased predictions of the individual cell populations in the mixed cell culture. Later Ram et al reiterated that “our approach was designed to predict growth in a mixed culture, with resource-based competition during a single growth phase, sampled at a high frequency” (Ram, Obolski, Feldman, Berman, & Hadany, 2020). Ram’s approach, presumably, may provide a convenient way to predict the growth fitness of a mixed cell population with resource competition. Our current work moves beyond resource competition and includes the social interactions of the mixed cell populations. Depending on the degree of freedom of the system, it might be possible to predict the community-level population dynamics from the biological parameter obtained in monoculture. Further experimental validation will be needed to corroborate this hypothesis.

The reported two species model is representative of microbial social interactions with two interaction coefficients *γ*_AB_ or *γ*_BA_. When we expand the model to include three species A, B and C, we could simply introduce six interaction coefficients, for example, *γ*_AB_, *γ*_AC_, *γ*_BA_, *γ*_BC_, *γ*_CA_ and *γ*_CA_. The growth fitness of one species (A) will be determined by two other species (B and C). The prediction of community behavior will be more complicated, possibly there will be multiple steady state solutions, interesting social structure or stratification may be emerged from multiple species interactions (*N* ≥3).

There are a number of studies have reported compelling cases to optimize microbial consortia performance by division of labor (Roell et al., 2019). Division of labor is especially useful to mitigate metabolic burden or metabolic stress when lengthy or incompatible pathways were expressed. Our current work deals with microbial social interaction and microbial competition, which are commonly found in naturally existing microbial species. To validate the reported experimental results, we need to introduce a stress factor to quantify the metabolic burden in consortia and correlate the burdensome effects with the growth fitness function. In addition, metabolite-host interactions, such as product or substrate inhibition, will be very complex, and the solution for such complicated system will inform us new knowledge and facilitate us to explore the optimal design criteria of synthetic microbial consortia. Our current model considers the ideal and simplified scenario: the stress factors and metabolite-host interactions have been lumped into the interaction coefficients *γ*_AB_ or *γ*_BA_, which describes the beneficial or detrimental interactions between species A and species B. Well-defined stress factors and metabolite-host interactions should be integrated to further expand the scope of this work.

## 5. Conclusions

Microbial consortium is a complex adaptive system with higher order dynamic characteristics that are not present by individual members. To accurately predict the social interactions, we formulate a set of unstructured kinetic models to quantitatively describe the dynamic interactions of multiple microbial species. With the generalized social interaction model (**Eqn.1** to **Eqn. 5**), we analytically derived the steady state solutions for the two interacting species and the substrate in the continuous stirred tank reactor (CSTR or chemostat). By computing the Jacobian matrix and evaluating the eigenvalues, we analyzed the stability of the possible co-existing states on the basis of eight social interaction categories: competition, coexisting parasitism, extinctive parasitism, cooperation, bistable amensalism, extinctive amensalism, coexisting commensalism and extinctive commensalism. Our model predicts that only parasitism, commensalism and cooperation could lead to stable co-existing state. We then move forward to understand the dynamics of microbial consortia with sequential metabolic reactions compartmentalized into distinct species. Coupled with Luedeking–Piret equation and Michaelis-Menten equation, accumulation of metabolic intermediate in one species and formation of final product in another species could be derived and assessed. We then conclude that there is inefficient conversion of metabolic intermediate to the final product if the two species form parasitism consortia. Our simulation indicates that commensalism consortia could efficiently convert metabolic intermediate to final product and maintain metabolic homeostasis (i.e., constant final product formation) with a broad range of dilution rates. Instead, cooperative consortia may not maintain this metabolic homeostasis due to the mutualistic relationship between the two species. In this study, we discovered the underlying dynamics and emergent properties of microbial consortia, which may provide critical knowledge for us to control and engineer multiple microbial species in a coculture system. In particular, the simplicity and the rich dynamics of the consortia model highlight the importance to integrate unstructured kinetic models and social interaction parameters to systematically improve our prediction power.

## Supporting information

Matlab codes

## Acknowledgments

Dr. Xu would like to acknowledge the Bill and Melinda Gates Foundation (OPP1188443) and National Science Foundation under grant number 1805139 for financially supporting this project.

## Conflicts of interests

None declared.

## Symbols and biophysical explanations with SI units

*μ*_Amax_: maximal specific growth rate for species A, (1/h)
*μ*_A_: specific growth rate for species A, (1/h)
*μ*_Bmax_: maximal specific growth rate for species B, (1/h)
*K*_B_: specific growth rate for species B, (1/h)
*K*_SA_: substrate saturation constant for species A, (g/L)
*K*_SB_: substrate saturation constant for species B, (g/L)
*Y*_AS_: Species A biomass yield from substrate S, (g/g)
*Y*_BS_: Species B biomass yield from substrate S, (g/g)
*Y*_BA_: Product B (*P*_B_) yield from intermediate A (*P*_A_), (g/g)
*Y*_PS_: Intermediate A (*P*_A_) yield from substrate S, (g/g)
*α*: Growth-associated intermediate A (*P*_A_) formation coefficient, dimensionless
*β*: Growth-unassociated intermediate A (*P*_A_) formation rate, (1/h)
*γ*_AB_: Interaction coefficient of species A imposes on species B, dimensionless
*γ*_BA_: Interaction coefficient of species B imposes on species A, dimensionless
*k*: Rate constant of intermediate A (*P*_A_) converted to product B (*P*_B_), (1/h)
*K*_*m*_: Intermediate A saturation constant for species B, (g/L)
*X*_*A*_: Species A biomass in the CSTR, (g/L)
*X*_*B*_: Species B biomass in the CSTR, (g/L)
*P*_A_: Intermediate A concentration in the CSTR, (g/L)
*P*_B_: Product B concentration in the CSTR, (g/L)
*S*: Substrate concentration in the CSTR, (g/L)
*S*_0_: Substrate concentration in the feeding stream, (g/L)
*D*: Dilution rate in the CSTR, (1/h)

## Tables

**Table 1.**
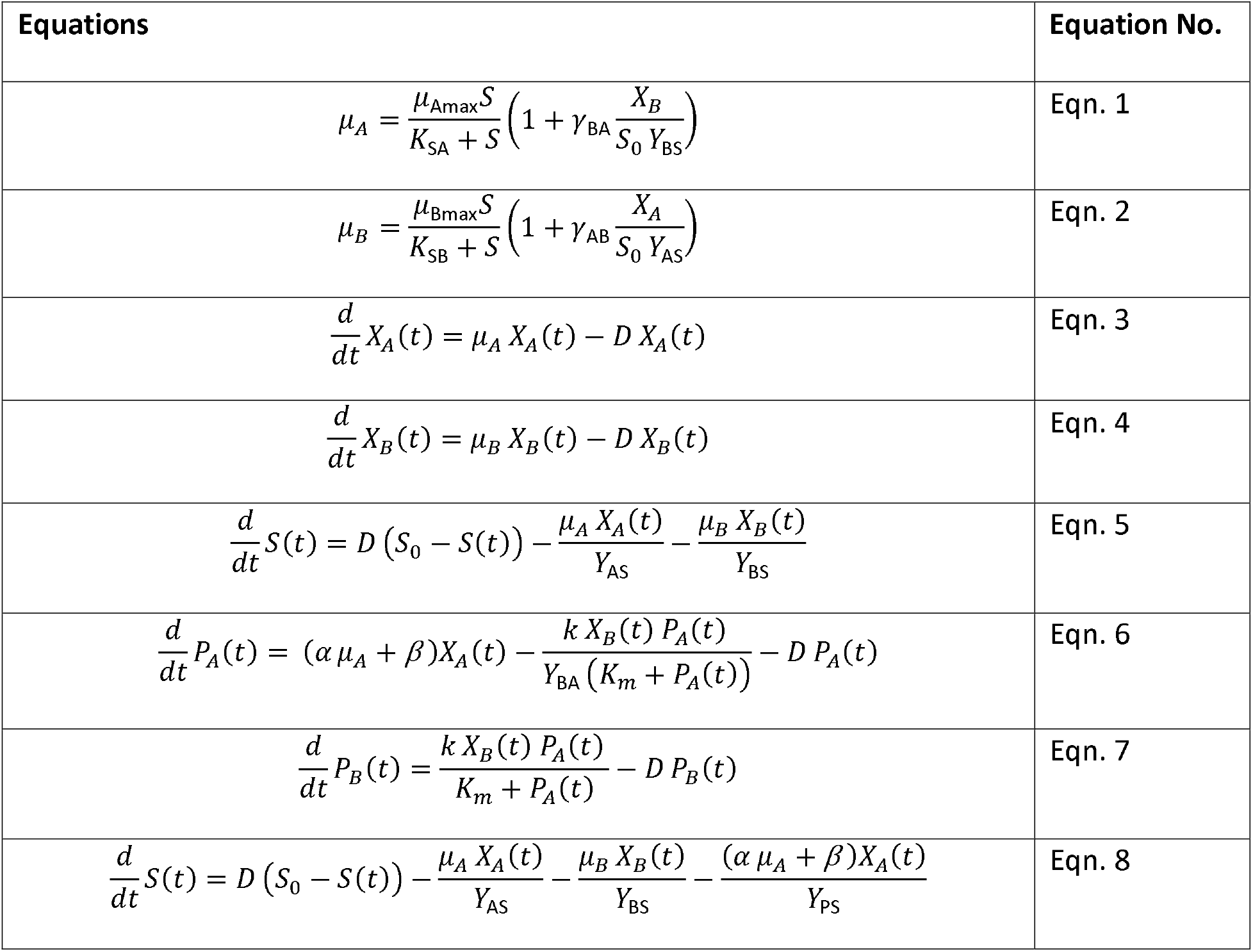
A unified mathematical model to describe the mass balance equations that govern microbial social interactions in chemostat.

| Equations | Equation No. |
| --- | --- |
| $\mu_A = \frac{\mu_{A\max} S}{K_{SA} + S} \left( 1 + \gamma_{BA} \frac{X_B}{S_0 Y_{BS}} \right)$ | Eqn. 1 |
| $\mu_B = \frac{\mu_{B\max} S}{K_{SB} + S} \left( 1 + \gamma_{AB} \frac{X_A}{S_0 Y_{AS}} \right)$ | Eqn. 2 |
| $\frac{d}{dt} X_A(t) = \mu_A X_A(t) - D X_A(t)$ | Eqn. 3 |
| $\frac{d}{dt} X_B(t) = \mu_B X_B(t) - D X_B(t)$ | Eqn. 4 |
| $\frac{d}{dt} S(t) = D (S_0 - S(t)) - \frac{\mu_A X_A(t)}{Y_{AS}} - \frac{\mu_B X_B(t)}{Y_{BS}}$ | Eqn. 5 |
| $\frac{d}{dt} P_A(t) = (\alpha \mu_A + \beta) X_A(t) - \frac{k X_B(t) P_A(t)}{Y_{BA} (K_m + P_A(t))} - D P_A(t)$ | Eqn. 6 |
| $\frac{d}{dt} P_B(t) = \frac{k X_B(t) P_A(t)}{K_m + P_A(t)} - D P_B(t)$ | Eqn. 7 |
| $\frac{d}{dt} S(t) = D (S_0 - S(t)) - \frac{\mu_A X_A(t)}{Y_{AS}} - \frac{\mu_B X_B(t)}{Y_{BS}} - \frac{(\alpha \mu_A + \beta) X_A(t)}{Y_{PS}}$ | Eqn. 8 |

## Notes

### Competing Interest Statement

The authors have declared no competing interest.

### Summary of Updates

Main text and supporting information changed

