## Supplementary material for "Dynamics of microbial competition, commensalism and cooperation and its implications for coculture and microbiome engineering": Matlab codes

Peng Xu*

Department of Chemical, Biochemical and Environmental Engineering, University of Maryland Baltimore County, Baltimore, MD 21250

#### 1. Supplementary Matlab code to generate the symbolic equations in this work

```
>> syms mu_A mu_B mu_Amax mu_Bmax K_SA K_SB S S0 Y_AS Y_BS X_A X_B gamma_AB
gamma_BA P_A P_B D alpha beta k K_m Y_BA Y_PS
>> eqn1 = mu_A == mu_Amax*S/(K_SA+S)*(1+gamma_BA*X_B/(S0*Y_BS));
>> eqn2 = mu_B == mu_Bmax*S/(K_SB+S)*(1+gamma_AB*X_A/(S0*Y_AS));
>> syms X_A(t) X_B(t) S(t) P_A(t) P_B(t)
>> eqn3 = diff(X_A,t) == mu_A*X_A-D*X_A;
>> eqn4 = diff(X_B,t) == mu_B*X_B-D*X_B;
>> eqn5 = diff(S,t) == D*(S0-S) -mu_A*X_A/Y_AS -mu_B*X_B/Y_BS;
>> eqn6 = diff(P_A,t) == (alpha*mu_A+beta)*X_A-D*P_A- k*X_B*P_A/(K_m+P_A)/Y_BA;
>> eqn7 = diff(P_B,t) == k*X_B*P_A/(K_m+P_A) -D*P_B;
>> eqn8 = diff(S,t) == D*(S0-S) -mu_A*X_A/Y_AS -mu_B*X_B/Y_BS -
(alpha*mu_A+beta)*X_A/Y_PS;
```

#### 2. Supplementary Matlab code to derive the analytical solution for Eqn1 to Eqn. 5

```
>> syms mu_A mu_B mu_Amax mu_Bmax K_SA K_SB S S0 Y_AS Y_BS X_A X_B gamma_AB
gamma_BA M1 M2 P1 P2 D alpha1 alpha2 beta1 beta2 Y_M1S Y_M2S Y_P1M1 Y_P2M2 k1 k2
K_M1 K_M2
>> eqn1 = mu_A == mu_Amax*S/(K_SA+S)*(1+gamma_BA*X_B/(S0*Y_BS));
>> eqn2 = mu_B == mu_Bmax*S/(K_SB+S)*(1+gamma_AB*X_A/(S0*Y_AS));
>> eqn3 = mu_A*X_A-D*X_A == 0;
>> eqn4 = mu_B*X_B-D*X_B ==0;
>> eqn5 = D*(S0-S) -mu_A*X_A/Y_AS -mu_B*X_B/Y_BS == 0;
```

```
>> eqns =[eqn1,eqn2, eqn3, eqn4, eqn5];
>> vars = [mu_A mu_B S X_A X_B];
>> [solmuA, solmuB, solS, solXA solXB] = solve(eqns, vars);
```

### 3. Supplementary Matlab code to derive the Jacobian matrix as defined by Eqn. 3-Eqn. 5

```
>> syms mu_Amax mu_Bmax K_SA K_SB S S0 Y_AS Y_BS X_A X_B gamma_AB gamma_BA D;

>> eqn1 = mu_Amax*S/(K_SA+S)*(1+gamma_BA*X_B/(S0*Y_BS))*X_A-D*X_A;

>> eqn2 = mu_Bmax*S/(K_SB+S)*(1+gamma_AB*X_A/(S0*Y_AS))*X_B-D*X_B;

>> eqn3 = D*(S0-S) - mu_Amax*S/(K_SA+S)*(1+gamma_BA*X_B/(S0*Y_BS))*X_A/Y_AS -
mu_Bmax*S/(K_SB+S)*(1+gamma_AB*X_A/(S0*Y_AS))*X_B/Y_BS;

>> JaM=jacobian([eqn1,eqn2,eqn3], [X_A,X_B,S])
```

$$Jacobian\ matrix\ =\ JaM\ =$$

$$\begin{pmatrix} \frac{S\,\mu_{\mathrm{Amax}}}{K_{\mathrm{SA}}+S}\left(\frac{X_B\,\gamma_{\mathrm{BA}}}{S_0\,Y_{\mathrm{BS}}}+1\right)-D & \frac{S\,X_A\,\gamma_{\mathrm{BA}}\,\mu_{\mathrm{Amax}}}{S_0\,Y_{\mathrm{BS}}\,(K_{\mathrm{SA}}+S)} & \frac{K_{\mathrm{SA}}\,X_A\,\mu_{\mathrm{Amax}}\,(S_0\,Y_{\mathrm{BS}}+X_B\,\gamma_{\mathrm{BA}})}{S_0\,Y_{\mathrm{BS}}\,(K_{\mathrm{SA}}+S)^2} \\ \frac{S\,X_B\,\gamma_{\mathrm{AB}}\,\mu_{\mathrm{Bmax}}}{S_0\,Y_{\mathrm{AS}}\,(K_{\mathrm{SB}}+S)} & \frac{S\,\mu_{\mathrm{Bmax}}}{K_{\mathrm{SB}}+S}\left(\frac{X_A\,\gamma_{\mathrm{AB}}}{S_0\,Y_{\mathrm{AS}}}+1\right)-D & \frac{K_{\mathrm{SB}}\,X_B\,\mu_{\mathrm{Bmax}}\,(S_0\,Y_{\mathrm{AS}}+X_A\,\gamma_{\mathrm{AB}})}{S_0\,Y_{\mathrm{AS}}\,(K_{\mathrm{SB}}+S)^2} \\ -\frac{S\,\mu_{\mathrm{Amax}}\left(\frac{X_B\,\gamma_{\mathrm{BA}}}{S_0\,Y_{\mathrm{BS}}}+1\right)}{Y_{\mathrm{AS}}\,(K_{\mathrm{SA}}+S)}-\frac{S\,X_B\,\gamma_{\mathrm{AB}}\,\mu_{\mathrm{Bmax}}}{S_0\,Y_{\mathrm{AS}}\,Y_{\mathrm{BS}}\,(K_{\mathrm{SB}}+S)} & -\frac{S\,\mu_{\mathrm{Bmax}}\left(\frac{X_A\,\gamma_{\mathrm{AB}}}{S_0\,Y_{\mathrm{AS}}}+1\right)}{Y_{\mathrm{BS}}\,(K_{\mathrm{SB}}+S)}-\frac{S\,X_A\,\gamma_{\mathrm{BA}}\,\mu_{\mathrm{Amax}}}{S_0\,Y_{\mathrm{AS}}\,Y_{\mathrm{BS}}\,(K_{\mathrm{SA}}+S)} & \frac{S\,X_A\,\mu_{\mathrm{Amax}}\left(\frac{X_B\,\gamma_{\mathrm{BA}}}{S_0\,Y_{\mathrm{BS}}}+1\right)}{Y_{\mathrm{AS}}\,(K_{\mathrm{SA}}+S)^2}-\frac{X_A\,\mu_{\mathrm{Amax}}\left(\frac{X_B\,\gamma_{\mathrm{BA}}}{S_0\,Y_{\mathrm{BS}}}+1\right)}{Y_{\mathrm{AS}}\,(K_{\mathrm{SA}}+S)}-\frac{X_B\,\mu_{\mathrm{Bmax}}\left(\frac{X_A\,\gamma_{\mathrm{AB}}}{S_0\,Y_{\mathrm{AS}}}+1\right)}{Y_{\mathrm{BS}}\,(K_{\mathrm{SB}}+S)}-D+\frac{S\,X_B\,\mu_{\mathrm{Bmax}}\left(\frac{X_A\,\gamma_{\mathrm{AB}}}{S_0\,Y_{\mathrm{AS}}}+1\right)}{Y_{\mathrm{BS}}\,(K_{\mathrm{SB}}+S)^2} \end{pmatrix}$$

Where we can simplify the Jacobian matrix by

$$J=\begin{pmatrix} A-D & C & AF \\ E & B-D & BG \\ -\frac{A}{Y_{\mathrm{AS}}}-\frac{E}{Y_{\mathrm{BS}}} & -\frac{B}{Y_{\mathrm{BS}}}-\frac{C}{Y_{\mathrm{AS}}} & -\frac{AF}{Y_{\mathrm{AS}}}-\frac{BG}{Y_{\mathrm{BS}}}-D \end{pmatrix}$$

$$A=\frac{S\,\mu_{\mathrm{Amax}}}{K_{\mathrm{SA}}+S}\left(\frac{X_B\,\gamma_{\mathrm{BA}}}{S_0\,Y_{\mathrm{BS}}}+1\right)$$

$$B=\frac{S\,\mu_{\mathrm{Bmax}}}{K_{\mathrm{SB}}+S}\left(\frac{X_A\,\gamma_{\mathrm{AB}}}{S_0\,Y_{\mathrm{AS}}}+1\right)$$

$$C=\frac{S\,X_A\,\gamma_{\mathrm{BA}}\,\mu_{\mathrm{Amax}}}{S_0\,Y_{\mathrm{BS}}\,(K_{\mathrm{SA}}+S)}$$

$$E=\frac{S\,X_B\,\gamma_{\mathrm{AB}}\,\mu_{\mathrm{Bmax}}}{S_0\,Y_{\mathrm{AS}}\,(K_{\mathrm{SB}}+S)}$$

$$F=\frac{K_{\mathrm{SA}}\,X_A}{(K_{\mathrm{SA}}+S)\,S}$$

$$G=\frac{K_{\mathrm{SB}}\,X_B}{(K_{\mathrm{SB}}+S)\,S}$$

### 4. Supplementary Matlab code to generate Fig. 6 in the manuscript

```
>> str1 = '#0072BD';
>> color1 = sscanf(str1(2:end),'%2x%2x%2x',[1 3])/255;
>> str2 = '#D95319';
>> color2 = sscanf(str2(2:end),'%2x%2x%2x',[1 3])/255;
>> str3 = '#7E2F8E';
>> color3 = sscanf(str3(2:end),'%2x%2x%2x',[1 3])/255;
>> str4 = '#77AC30';
>> color4 = sscanf(str4(2:end),'%2x%2x%2x',[1 3])/255;
>> str5 = '#4DBEEE';
>> color5 = sscanf(str5(2:end),'%2x%2x%2x',[1 3])/255;

>> subplot(2,2,1)
>> for D = 0.06:0.0025:2.0;
>> tspan = 0:0.25:500;
>> [T,Y]= ode45(@(t,y) coculture13(t,y,D),tspan, [1 1 1 1 1]);
>> yyaxis left;
>> plot(D, Y(2000,2), '.', 'Color', color1);
>> hold on;
>> plot(D, Y(2000,4), '.', 'Color', color3);
>> plot(D, Y(2000,5), '.', 'Color', color4);
>> yyaxis right;
>> plot(D, Y(2000,1), '.', 'Color', color2);
>> plot(D, Y(2000,3), '.', 'Color', color5);
>> hold on;
>> end
>> xlabel(' D (1/h)');
>> xlim([0 1.8]);
>> yyaxis left;
>> ylabel('X_A, P_A, P_B (g/L)');
>> ylim([0 10]);
>> yyaxis right;
>> ylabel('S, X_B (g/L)');
>> ylim([0 50]);
>> grid on;
>> title('(a) \gamma_{AB} = +1; \gamma_{BA} = -1');
>> hold off;

>> subplot(2,2,2);
>> for D = 0.01:0.005:2.0;
>> tspan = 0:0.25:500;
>> [T,Y]= ode45(@(t,y) coculture15(t,y,D),tspan, [1 1 1 1 1]);
>> yyaxis left;
>> plot(D, Y(2000,2), '.', 'Color', color1);
>> hold on;
>> plot(D, Y(2000,3), '.', 'Color', color5);
>> plot(D, Y(2000,4), '.', 'Color', color3);
>> plot(D, Y(2000,5), '.', 'Color', color4);
>> yyaxis right;
>> plot(D, Y(2000,1), '.', 'Color', color2);
>> hold on;
```

```
>> end
>> xlabel(' D (1/h)');
>> xlim([0 1.8]);
>> yyaxis left;
>> ylabel('X_A, X_B, P_A, P_B (g/L)');
>> ylim([0 20]);
>> yyaxis right;
>> ylabel('S (g/L)');
>> ylim([0 50]);
>> grid on;
>> title('(b) \gamma_{AB} = -1; \gamma_{BA} = +1');
>> hold off;

>> subplot(2,2,3);
>> for D = 0.01:0.005:2.0;
>> tspan = 0:0.25:500;
>> [T,Y]= ode45(@(t,y) coculture12(t,y,D),tspan, [1 1 1 1 1]);
>> yyaxis left;
>> plot(D, Y(2000,2), '.', 'Color', color1);
>> hold on;
>> plot(D, Y(2000,3), '.', 'Color', color5);
>> plot(D, Y(2000,4), '.', 'Color', color3);
>> plot(D, Y(2000,5), '.', 'Color', color4);
>> yyaxis right;
>> plot(D, Y(2000,1), '.', 'Color', color2);
>> hold on;
>> end
>> xlabel(' D (1/h)');
>> xlim([0 1.8]);
>> yyaxis left;
>> ylabel('X_A, X_B, P_A, P_B (g/L)');
>> ylim([0 25]);
>> yyaxis right;
>> ylabel('S (g/L)');
>> ylim([0 50]);
>> Legend=cell(4,1)
>> Legend{1}='X_A' ;
>>  Legend{2}='X_B';
>>  Legend{3}='P_A' ;
>>  Legend{4}='P_B';
>>  legend(Legend);
>> legend boxoff
>> grid on;
>> title('(c) \gamma_{AB} = +1; \gamma_{BA} = 0');
>> hold off;

>> subplot(2,2,4);
>> for D = 0.01:0.0025:2.0;
>> tspan = 0:0.25:500;
>> [T,Y]= ode45(@(t,y) coculture11(t,y,D),tspan, [1 1 1 1 1]);
>> yyaxis left;
>> plot(D, Y(2000,2), '.', 'Color', color1);
```

```
>> hold on;
>> plot(D, Y(2000,3), '.', 'Color', color5);
>> plot(D, Y(2000,4), '.', 'Color', color3);
>> plot(D, Y(2000,5), '.', 'Color', color4);
>> yyaxis right;
>> plot(D, Y(2000,1), '.', 'Color', color2);
>> hold on;
>> end
>> xlabel(' D (1/h)');
>> xlim([0 1.8]);
>> yyaxis left;
>> ylabel('X_A, X_B, P_A, P_B (g/L)');
>> ylim([0 16]);
>> yyaxis right;
>> ylabel('S (g/L)');
>> ylim([0 50]);
>> Legend=cell(4,1)
>>  Legend{1}='X_A' ;
>>  Legend{2}='X_B';
>>  Legend{3}='P_A' ;
>>  Legend{4}='P_B';
>>  legend(Legend);
>> legend boxoff
>> grid on;
>> title('(d) \gamma_{AB} = +1; \gamma_{BA} = +1')
```

### 5. Representative m. files to generate the analytical solutions and the trajectory

**m.file # 1**

```
function dydt = coculture1(t,y,D)
dydt=zeros(3,1);

mu_Amax=1.6; mu_Bmax=1.2; K_SA=1.0; K_SB=0.8; S0=50; Y_AS=0.5; Y_BS=0.8;
gamma_BA=-1;gamma_AB=-1;

S   = y(1);
X_A = y(2);
X_B = y(3);

% Define the two specific growth rate
mu_A = mu_Amax*S/(K_SA+S)*(1+gamma_BA*X_B/(S0*Y_BS));
mu_B = mu_Bmax*S/(K_SB+S)*(1+gamma_AB*X_A/(S0*Y_AS));

dydt(1) = D*(S0-S) -mu_A*X_A/Y_AS -mu_B*X_B/Y_BS;
dydt(2) = mu_A*X_A-D*X_A;
dydt(3) = mu_B*X_B-D*X_B;

end
```

**m.file # 2**

```
function dydt = coculture2(t,y,D)
dydt=zeros(3,1);


mu_Amax=1.6; mu_Bmax=1.2; K_SA=1.0; K_SB=0.8; S0=50; Y_AS=0.5; Y_BS=0.8;
gamma_BA=-1;gamma_AB=0;

S   = y(1);
X_A = y(2);
X_B = y(3);

% Define the two specific growth rate
mu_A = mu_Amax*S/(K_SA+S)*(1+gamma_BA*X_B/(S0*Y_BS));
mu_B = mu_Bmax*S/(K_SB+S)*(1+gamma_AB*X_A/(S0*Y_AS));

dydt(1) = D*(S0-S) -mu_A*X_A/Y_AS -mu_B*X_B/Y_BS;
dydt(2) = mu_A*X_A-D*X_A;
dydt(3) = mu_B*X_B-D*X_B;


end
```

**m.file # 3**

```
function dydt = coculture3(t,y,D)
dydt=zeros(3,1);

mu_Amax=1.6; mu_Bmax=1.2; K_SA=1.0; K_SB=0.8; S0=50; Y_AS=0.5; Y_BS=0.8;
gamma_BA=-1;gamma_AB=1;

S   = y(1);
X_A = y(2);
X_B = y(3);


% Define the two specific growth rate
mu_A = mu_Amax*S/(K_SA+S)*(1+gamma_BA*X_B/(S0*Y_BS));
mu_B = mu_Bmax*S/(K_SB+S)*(1+gamma_AB*X_A/(S0*Y_AS));

dydt(1) = D*(S0-S) -mu_A*X_A/Y_AS -mu_B*X_B/Y_BS;
dydt(2) = mu_A*X_A-D*X_A;
dydt(3) = mu_B*X_B-D*X_B;

end
```

**m.file # 4**

```
function dydt = coculture4(t,y,D)
dydt=zeros(3,1);


mu_Amax=1.6; mu_Bmax=1.2; K_SA=1.0; K_SB=0.8; S0=50; Y_AS=0.5; Y_BS=0.8;
gamma_BA=0;gamma_AB=-1;

S   = y(1);
X_A = y(2);
X_B = y(3);


% Define the two specific growth rate
mu_A = mu_Amax*S/(K_SA+S)*(1+gamma_BA*X_B/(S0*Y_BS));
mu_B = mu_Bmax*S/(K_SB+S)*(1+gamma_AB*X_A/(S0*Y_AS));

dydt(1) = D*(S0-S) -mu_A*X_A/Y_AS -mu_B*X_B/Y_BS;
dydt(2) = mu_A*X_A-D*X_A;
dydt(3) = mu_B*X_B-D*X_B;


end
```

**m.file # 16**

```
function dydt = coculture16(t,y,D)
dydt=zeros(5,1);

mu_Amax=1.6; mu_Bmax=1.2; K_SA=1.0; K_SB=0.8; S0=50; Y_AS=0.5; Y_BS=0.8;
gamma_BA=-1;gamma_AB=0;alpha=0.5;beta=0.5;Y_PS=0.4;Y_BA=0.8; k=0.8; K_m = 1.0;

S   = y(1);
X_A = y(2);
X_B = y(3);
P_A = y(4);
P_B = y(5);

% Define the two specific growth rate
mu_A = mu_Amax*S/(K_SA+S)*(1+gamma_BA*X_B/(S0*Y_BS));
mu_B = mu_Bmax*S/(K_SB+S)*(1+gamma_AB*X_A/(S0*Y_AS));

dydt(1) = D*(S0-S) -mu_A*X_A/Y_AS -mu_B*X_B/Y_BS - (alpha*mu_A+beta)*X_A/Y_PS;
dydt(2) = mu_A*X_A-D*X_A;
dydt(3) = mu_B*X_B-D*X_B;
dydt(4) = (alpha*mu_A+beta)*X_A-D*P_A- k*X_B*P_A/(K_m+P_A)/Y_BA;
dydt(5) = k*X_B*P_A/(K_m+P_A) -D*P_B;

end
```

### 6. Supplementary Tables: representative eigenvalues computed in this study

| Representative social interactions | Dilution rate (1/h) | Steady state solutions | | | Three eigenvalues with respect to each steady state | | | Category | Stable or Unstable |
|---|---|---|---|---|---|---|---|---|---|
| | | $S$ (g/L) | $X_A$ (g/L) | $X_B$ (g/L) | $\lambda_1$ | $\lambda_2$ | $\lambda_3$ | | |
| Competition $\gamma_{AB} = -1$ $\gamma_{BA} = -1$ | $D$ = 1.0 | 50.000 | 0.000 | 0.000 | -1.00 | 0.18 | 0.57 | washout | Unstable |
| | | 1.667 | 24.167 | 0.000 | -10.88 | -1.00 | -0.97 | A survive | Stable |
| | | 4.000 | 0.000 | 36.800 | -1.92 | -1.00 | -0.90 | B survive | Stable |
| | | 25.454 | 3.512 | 14.018 | -1.00 | 0.30 | -0.33 | **Coexist** | Unstable |
| | | -2.537 | 10.735 | 24.853 | -1.00 | -12.25 | 1.11 | Nonsense | Unstable |
| Amensalism $\gamma_{AB} = 0$ $\gamma_{BA} = -1$ | $D$ = 1.0 | 50.000 | 0.000 | 0.000 | -1.00 | 0.18 | 0.57 | washout | Unstable |
| | | 1.667 | 24.167 | 0.000 | -10.88 | -1.00 | -0.19 | A survive | Stable |
| | | 4.000 | 0.000 | 36.800 | -1.92 | -1.00 | -0.90 | B survive | Stable |
| | | 4.000 | 17.531 | 8.750 | -1.00 | -2.38 | 0.17 | **Coexist** | Unstable |
| Parasitism $\gamma_{AB} = +1$ $\gamma_{BA} = -1$ | $D$ = 1.0 | 50.00 | 0.00 | 0.00 | -1.00 | 0.18 | 0.57 | washout | Unstable |
| | | 1.67 | 24.17 | 0.00 | -10.88 | -1.00 | 0.59 | A survive | Unstable |
| | | 4.00 | 0.00 | 36.80 | -1.92 | -1.00 | -0.90 | B survive | Stable |
| | | 0.05 | 312.08 | -459.4 | -1.00 | -1027.61 | -0.62 | Nonsense | Stable |
| | | 39.53 | -3.75 | 14.37 | -1.00 | 0.31 | -0.32 | Nonsense | Unstable |
| Parasitism $\gamma_{AB} = +1$ $\gamma_{BA} = -1$ | $D$ = 1.365 | 50.00 | 0.00 | 0.00 | -1.37 | -0.18 | 0.20 | washout | Unstable |
| | | 5.81 | 22.10 | 0.00 | -1.53 | -1.37 | 0.62 | A survive | Unstable |
| | | -6.62 | 0.00 | 45.29 | -1.61 | -1.61 | -1.37 | Nonsense | Stable |
| | | 0.08 | 289.05 | -422.5 | -1.37 | -0.74 | -943.44 | Nonsense | Stable |
| | | 35.70 | 4.07 | 4.92 | -1.37 | **- 0.0068 + 0.19*i*** | **-0.0068 - 0.19*i*** | **Coexist** | **Stable** |
| Amensalism $\gamma_{AB} = -1$ $\gamma_{BA} = 0$ | $D$ = 1.0 | 50.00 | 0.00 | 0.00 | -1.00 | 0.18 | 0.57 | Washout | Unstable |
| | | 1.67 | -5.83 | 48.00 | -1.00 | -8.76 | -0.29 | Nonsense | Stable |
| | | 1.67 | 24.17 | 0.00 | -10.88 | -1.00 | -0.97 | A survive | Stable |
| | | 4.00 | 0.00 | 36.80 | -1.92 | -1.00 | 0.28 | B survive | Unstable |
| Neutralism $\gamma_{AB} = 0$ $\gamma_{BA} = 0$ | $D$ = 1.0 | 50.00 | 0.00 | 0.00 | -1.00 | 0.18 | 0.57 | washout | Unstable |
| | | 1.67 | 24.17 | 0.00 | -10.88 | -1.00 | -0.19 | A survive | Stable |
| | | 4.00 | 0.00 | 36.80 | -1.92 | -1.00 | 0.28 | B survive | Unstable |
| Commensalism $\gamma_{AB} = +1$ $\gamma_{BA} = 0$ | $D$ = 1.0 | 50.000 | 0.000 | 0.000 | -1.00 | 0.18 | 0.57 | Washout | Unstable |
| | | 1.667 | 5.833 | 29.333 | -1.00 | -9.60 | -0.16 | **Coexist** | **Stable** |
| | | 1.667 | 24.167 | 0.000 | -10.88 | -1.00 | 0.59 | A survive | Unstable |

| | | | | | | | | | |
|---|---|---|---|---|---|---|---|---|---|
| | | 4.000 | 0.000 | 36.800 | -1.92 | -1.00 | 0.28 | B survive | Unstable |
| Parasitism<br>$\gamma_{\mathrm{AB}} = -1$<br>$\gamma_{BA} = +1$ | $D$ = 1.0 | 50.00 | 0.00 | 0.00 | -1.00 | 0.18 | 0.57 | washout | Unstable |
| | | 1.67 | 24.17 | 0.00 | -10.88 | -1.00 | -0.97 | A survive | Stable |
| | | 4.00 | 0.00 | 36.80 | -1.92 | -1.00 | 1.46 | B survive | Unstable |
| | | -0.03 | 487.78 | -740.4 | -1.00 | -1.56 | 1254.78 | Nonsense | Unstable |
| | | 60.45 | 3.89 | -14.59 | -1.00 | 0.18 | 0.57 | Nonsense | Unstable |
| Commensalism<br>$\gamma_{\mathrm{AB}} = 0$<br>$\gamma_{BA} = +1$ | $D$ = 1.0 | 50.00 | 0.00 | 0.00 | -1.00 | 0.18 | 0.57 | Washout | Unstable |
| | | 1.67 | 24.17 | 0.00 | -10.88 | -1.00 | -0.19 | A survive | Stable |
| | | 4.00 | 0.00 | 36.80 | -1.92 | -1.00 | 1.46 | B survive | Unstable |
| | | 4.00 | 28.47 | -8.75 | -1.00 | -2.64 | 0.25 | Nonsense | Unstable |
| Cooperation<br>$\gamma_{\mathrm{AB}} = +1$<br>$\gamma_{BA} = +1$ | $D$ = 1.0 | 50.00 | 0.00 | 0.00 | -1.00 | 0.18 | 0.57 | Washout | Unstable |
| | | 1.67 | 24.17 | 0.00 | -10.88 | -1.00 | 0.59 | A survive | Unstable |
| | | 4.00 | 0.00 | 36.80 | -1.92 | -1.00 | 1.46 | B survive | Unstable |
| | | 76.24 | -3.95 | -14.67 | -1.00 | -0.33 | 0.33 | Nonsense | Unstable |
| | | 0.85 | 15.51 | 14.51 | -1.00 | -29.92 | -0.30 | **Coexist** | **Stable** |
| Cooperation<br>$\gamma_{\mathrm{AB}} = +1$<br>$\gamma_{BA} = +1$ | $D$ =<br>1.365 | 50.00 | 0.00 | 0.00 | -1.37 | -0.18 | 0.20 | Washout | Unstable |
| | | 5.81 | 22.10 | 0.00 | -1.53 | -1.37 | 0.62 | A survive | Unstable |
| | | -6.62 | 0.00 | 45.29 | -1.61 | -1.37 | 2.65 | Nonsense | Unstable |
| | | 48.66 | 3.91 | -5.17 | -1.37 | **- 0.0007<br>+ 0.19*i*** | **-0.0007<br>- 0.19*i*** | Nonsense | Stable |
| | | 1.81 | 15.99 | 12.96 | -1.37 | -0.39 | -11.92 | **Coexist** | **Stable** |
| Cooperation<br>$\gamma_{\mathrm{AB}} = +1$<br>$\gamma_{BA} = +1$ | $D$ =<br>1.60 | 50.00 | 0.00 | 0.00 | -1.60 | -0.42 | -0.03 | **Washout** | Stable |
| | | -3.20 | 0.00 | 42.56 | -8.87 | -1.60 | 3.20 | Nonsense | Unstable |
| | | 29.87 | 9.23 | 1.34 | -1.60 | 0.12 | -0.16 | **Coexist** | Unstable |
| | | 3.46 | 16.04 | 11.56 | -1.60 | -4.17 | -0.41 | **Coexist** | **Stable** |
| Cooperation<br>$\gamma_{\mathrm{AB}} = +1$<br>$\gamma_{BA} = +1$ | $D$ =<br>1.70 | 50.00 | 0.00 | 0.00 | -1.700 | -0.519 | -0.131 | **Washout** | Stable |
| | | -17.00 | 33.50 | 0.00 | -1.700 | -0.419 | 1.247 | Nonsense | Unstable |
| | | -2.72 | 0.00 | 42.18 | -13.729 | -1.700 | 3.498 | Nonsense | Unstable |
| | | 20.75 | 11.78 | 4.55 | -1.700 | 0.218 | -0.324 | **Coexist** | Unstable |
| | | 5.29 | 15.77 | 10.53 | -1.700 | -0.381 | -1.785 | **Coexist** | **Stable** |
